# SQANTI-browser: visualization and curation of SQANTI3-classified long-read transcriptomes within the UCSC Genome Browser

**DOI:** 10.64898/2026.05.25.727625

**Authors:** Alejandro Paniagua, Carlos Blanco, Andrés Colomer Fernández, Mark Diekhans, Ana Conesa, Carolina Monzó

## Abstract

Long-read sequencing enables transcriptome-wide isoform discovery. However, it generates substantial technical and structural ambiguity that complicates transcript interpretation. Here, we present SQANTI-browser, a classification-aware visualization framework that converts SQANTI3 outputs into interactive UCSC Genome Browser Track Hubs, preserving full transcript structural metadata. By integrating SQANTI classifications directly within the UCSC ecosystem, SQANTI-browser enables dynamic filtering and evidence-guided curation alongside public resource tracks. Furthermore, its adaptive architecture natively supports non-reference genomes, orthogonal data, and custom metadata fields. Applied to clinical, noisy, and synthetic datasets, SQANTI-browser resolves alignment artifacts and rescues actionable novel isoforms, providing a robust framework for long-read transcriptome curation.

## Background

Long-read sequencing (LRS) technologies from Pacific Biosciences and Oxford Nanopore Technologies enable direct isoform-resolved transcriptome profiling by capturing full-length RNA molecules from transcription start to termination sites [1]. The widespread adoption of LRS has revealed extensive transcriptomic complexity across tissues and species, including large numbers of transcripts that differ from existing annotations [2–4]. As reconstruction algorithms improve, the analytical challenge has shifted from isoform discovery to the rigorous evaluation of structural novelty and technical artifacts [5, 6].

The SQANTI3 framework provides systematic structural classification and quality control for long-read transcriptomes by assigning transcript models to standardized categories and computing quantitative descriptors of splice junction support, coding potential, and positional agreement with reference annotations [7]. While these outputs enable reproducible filtering and benchmarking, they are primarily distributed as tabular files and static annotations. Consequently, the evidence supporting each transcript model is separated from its genomic context, complicating expert inspection and iterative curation.

Effective interpretation of long-read transcriptomes requires simultaneous visualization of exon–intron structure, classification status, and quantitative quality metrics within a unified genomic coordinate system. General-purpose genome browsers such as Integrative Genomics Viewer [8], JBrowse [9] and the UCSC Genome Browser [10] support flexible track display but do not natively preserve structured long-read classification schemas as queryable attributes. As a result, evaluation of candidate novel isoforms often requires manual cross-referencing between browser views and external classification tables.

Here we present SQANTI-browser, a visualization and curation tool that converts SQANTI3 outputs into classification-aware Track Hubs compatible with both the UCSC Genome Browser [10, 11] and the Ensembl browser [12]. SQANTI-browser provides an automated pipeline to encode the complete SQANTI3 attribute space directly within indexed browser tracks using an extended AutoSQL schema, enabling transcript models to retain their structural categories and associated quality metrics within the visualization layer. This design allows dynamic filtering and attribute-based querying of isoforms directly in the browser interface, without relying on external data manipulation or command-line scripting. By embedding classification metadata within standard UCSC infrastructure, SQANTI-browser enables integrated inspection of long-read transcript models together with reference annotations and public genomic datasets in a single, reproducible environment.

We demonstrate the utility of SQANTI-browser across three representative use cases: discovery of a clinically relevant isoform, interactive curation of noisy loci, and resolution of splice-junction discrepancies in synthetic controls.

## Results

### Overview of the SQANTI-browser framework

SQANTI-browser converts SQANTI3 transcript classification outputs into a UCSC-compatible Track Hub that preserves the complete structural and quality metadata associated with each isoform (Fig. 1). The software requires as input the curated transcriptome annotation (.gtf) and the corresponding SQANTI3 classification file. Optional auxiliary files (e.g., junction support tables, CAGE and PolyA peaks) can also be incorporated to extend metadata fields.

**Figure 1.**
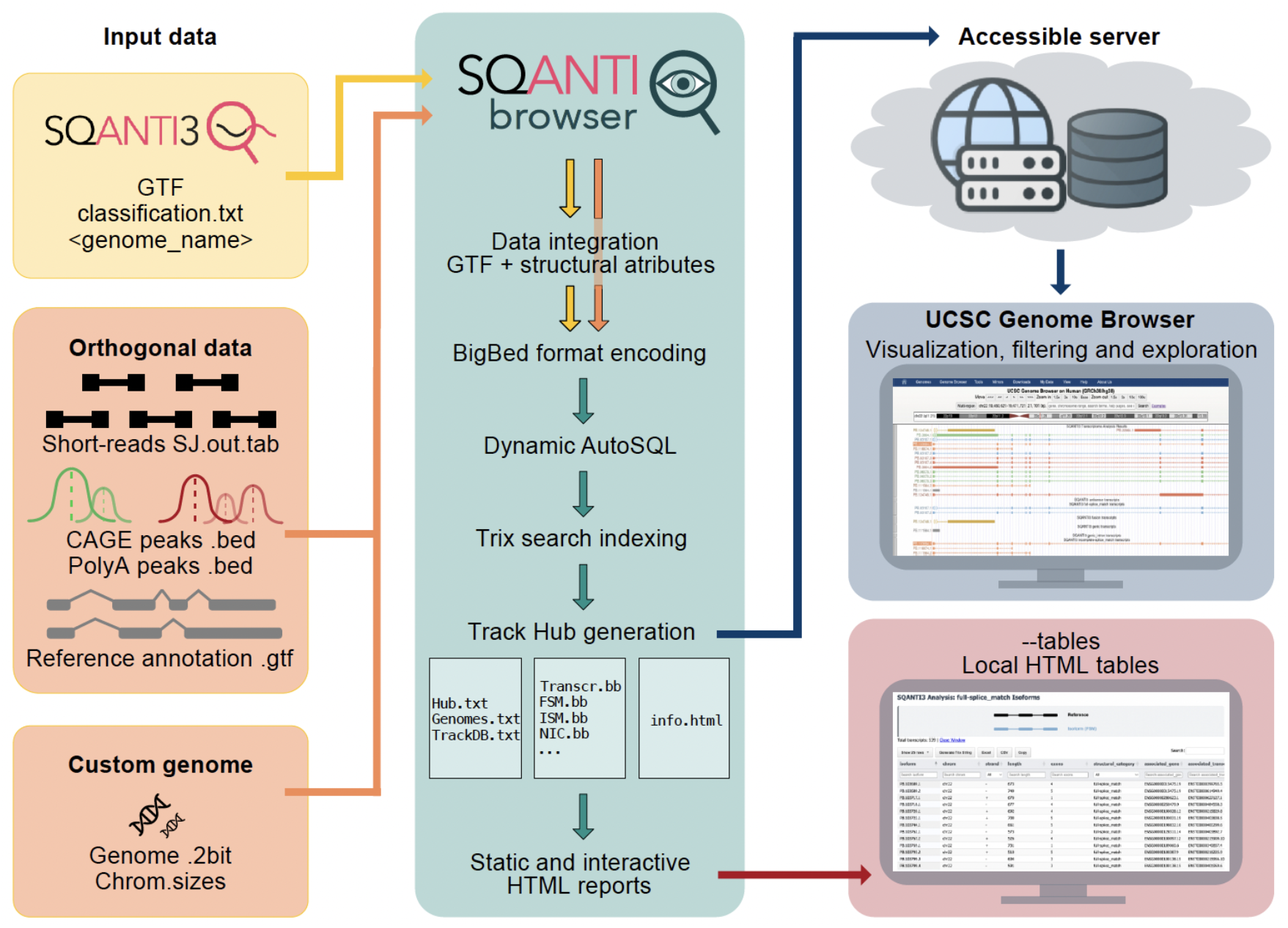
The SQANTI-browser framework for interactive long-read transcriptome curation. SQANTI-browser integrates SQANTI3 classification outputs with optional orthogonal validation data, including CAGE-seq peaks, polyA sites, and short-read splice junctions. The core engine processes transcript models and quality metrics into binary indexed formats (bigBed) using a dynamically generated AutoSQL schema for metadata-rich visualization. A Trix index is built to support multi-attribute searching across the dataset. The workflow generates a complete UCSC-compatible Track Hub, featuring whole-transcriptome and category-specific tracks (full-splice match: FSM, incomplete-splice match: ISM, novel in catalog: NIC, novel not in catalog: NNC, etc.), and interactive HTML reports for multidimensional filtering and quality assessment.

During processing, SQANTI-browser performs the following steps:

1. Metadata compatibilization. Transcript coordinates, structural categories, and quality descriptors are parsed from the SQANTI3 .gtf and classification files and standardized into a unified attribute table compatible with BED12+ UCSC Genome Browser formats.
2. Schema definition. An adaptive AutoSQL definition is automatically constructed to accommodate all metadata attributes detected.
3. Track compilation. Transcript models are converted into binary indexed bigBed files, embedding both exon–intron structures and all associated metadata fields.
4. Track Hub assembly. The required hub.txt, genomes.txt, and trackDb.txt configuration files are generated automatically, together with optional Trix indices to enable keyword-based search within the UCSC Genome Browser. Optional companion bigBeds for validation overlays (CAGE, polyA etc) are registered in the same hub.
5. Interactive reporting. Optionally, the software generates local interactive HTML reports providing a searchable, tabular summary of the transcriptome. These reports facilitate the exploration and export of curated subsets, while generating custom search strings to instantly locate specific isoforms within the UCSC interface.

The resulting Track Hub can be deployed on any static web server supporting byte-range requests. Once hosted, the transcriptome is accessible through a single URL and can be loaded directly into the UCSC Genome Browser without additional configuration.

Within the browser interface, each isoform retains all SQANTI3 annotations, including structural category, splice junction support metrics, coding potential scores, and positional annotations relative to reference transcripts. These attributes can be used for dynamic filtering and selective visualization directly in the UCSC display panel. Because the output conforms to the Track Hub specification, SQANTI-browser tracks can be visualized alongside all UCSC publicly available datasets, allowing structural classifications to be interpreted in the context of independent experimental evidence, such as epigenetic marks [13, 14], evolutionary conservation [15], or population-scale variation [16].

SQANTI-browser is maintained in parallel with SQANTI3 [7], SQANTI-reads [17] and other SQANTI-verse tools to ensure compatibility with evolving classification outputs and metadata structures.

### Discovering a high-confidence novel isoform in a clinically relevant locus

To demonstrate the utility of SQANTI-browser in a discovery-driven context, we analyzed the isoform-centric microglia genomic atlas, a resource of long-read transcriptomes derived from 30 post-mortem human brains [18]. First, we annotated the atlas using the SQANTI classification nomenclature [7, 19], where the main transcript structural categories are full-splice match (FSM) when the transcripts have the same exons and internal junction chain as the reference; incomplete-splice match (ISM) when they have fewer 5’ and/or 3’ exons than the reference, but their internal junctions match the reference annotation; novel in catalog (NIC) when they are not found in the reference, but carry a combination of known donor/acceptor sites; and novel not in catalog (NNC) when they carry at least one donor or acceptor site that is not annotated in the reference. Next, using the SQANTI-browser interactive HTML table, we applied classification-aware filtering to identify high-confidence novel isoforms. Specifically, we prioritized transcripts that were predicted as coding [20, 21], exhibited high positional concordance (+/-50 bp) with reference FANTOM5 CAGE peaks (transcription start site; TSS) [13] and PolyASite Atlas (transcription termination site; TTS) [22] entries, and excluded transcripts flagged for potential technical artifacts such as reverse transcriptase template switching (RTS), predicted nonsense-mediated decay (NMD) or non-canonical splice sites (Fig. 2a, Supplementary File 1).

**Figure 2.**
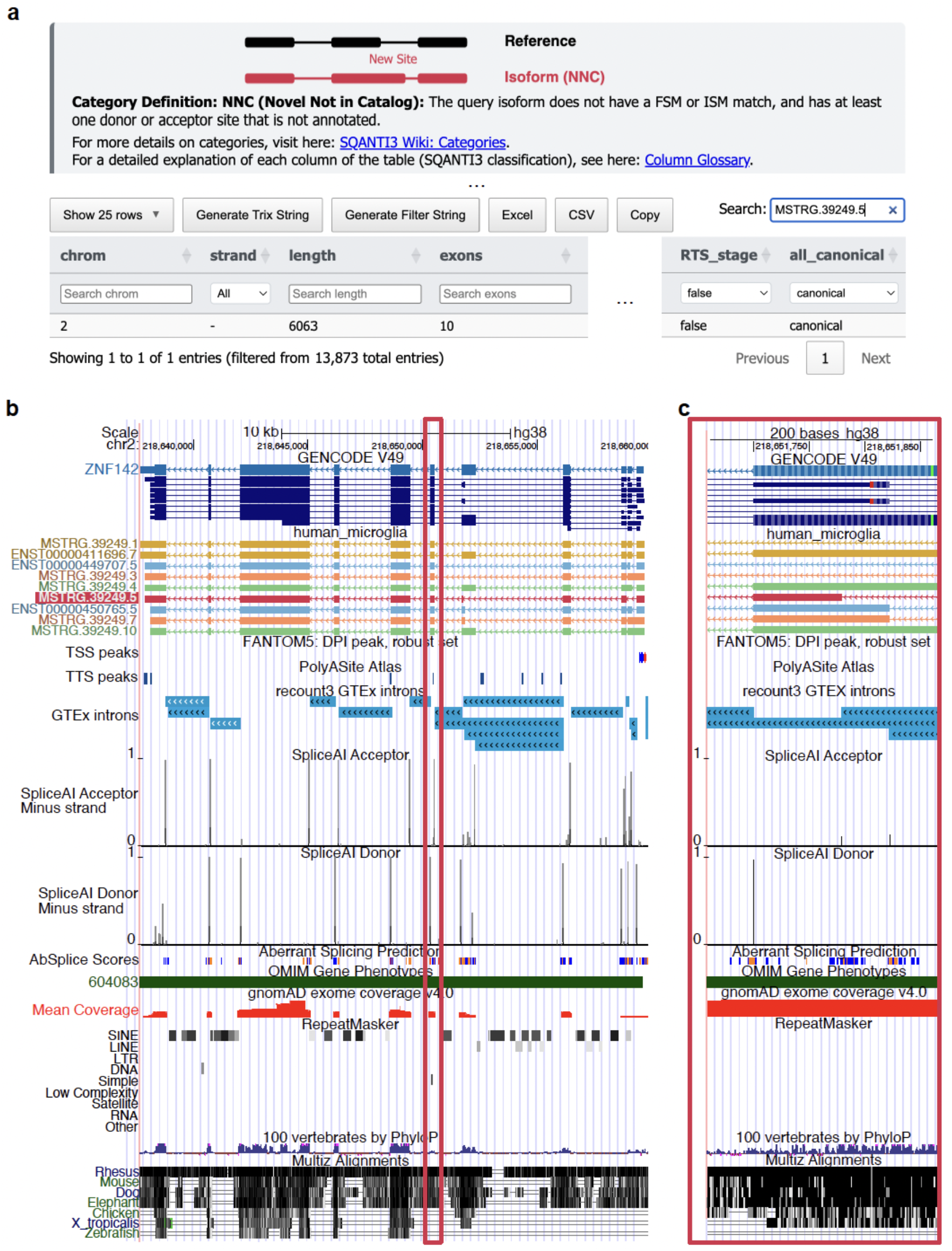
High-confidence novel isoform in a clinically relevant locus. **a** SQANTI-browser’s HTML table with filtered isoforms. **b** SQANTI-browser visualization of the complete *ZNF142* gene including orthogonal data tracks. **c** Zoomed SQANTI-browser visualization of the curated Novel Not in Catalog (NNC: red; MSTRG.39249.5) exon identified in the *ZNF142* locus.

This strategy identified a novel isoform of *ZNF142*, a zinc-finger protein gene associated with an autosomal recessive neurodevelopmental disorder characterized by impaired intellectual development and hyperkinetic movements (OMIM: 618425) [23]. Trix-based search (Supplementary Fig. 1) and visual inspection in the UCSC Genome Browser revealed that the NNC classification of this transcript arises from a shortened exon, generated by the usage of an alternative, unannotated splice site that produces a distinct exon boundary relative to the GENCODE reference [2] (Fig. 2b).

The existence of this novel junction was independently corroborated by additional databases available within the UCSC Genome Browser. Orthogonal short-read coverage from the GTEx project [24, 25] indicated clear splice-junction support at the alternative boundary. Furthermore, predictive deep-learning models (SpliceAI [26] and AbSplice [27]) assigned high probability scores to the novel splice site (Fig. 2b). In addition, the alternative exon boundary lies within a region of evolutionary conservation across vertebrates (UCSC 100 Vertebrates [15]) and is well covered by population-scale sequencing data (gnomAD [16]), supporting its genomic stability and functional plausibility. Finally, the absence of nearby repetitive elements in the RepeatMasker [28] track further argues against repeat-driven splicing artifacts or mapping errors.

Crucially, interactive evaluation of the transcript’s coding potential within the SQANTI-browser UCSC framework further supported the biological plausibility of the isoform. The novel transcript achieved a near-perfect Psauron score (0.98), while TransDecoder2 predicted a complete coding sequence (CDS) of 4,575 nucleotides that preserved the reading frame, yielding a predicted protein of 1,525 amino acids. Together, these analyses indicate that usage of the alternative splice site is compatible with maintenance of an intact open reading frame rather than generating a truncated or disrupted transcript product. The ability to simultaneously integrate microglial long-reads with predictive splicing models, population-scale databases, and ORF annotations allowed us to prioritize this NNC as a high-confidence candidate coding isoform, demonstrating how classification-aware integrated visualization facilitates the identification of biologically relevant transcript diversity absent from standard annotations like GENCODE [2] or RefSeq [29].

### Interactively curating candidate isoforms in noisy long-read transcriptomes

While high-confidence prioritization can identify biologically relevant isoforms in clean datasets, long-read transcriptomes frequently contain loci where technical artifacts and biological heterogeneity complicate transcriptome interpretation [6]. We used SQANTI-browser to iteratively inspect and curate transcript models across three challenging scenarios: degradation-associated sequence fragmentation, cross-platform technical bias, and age-related transcriptomic variation.

Partially degraded RNA molecules characteristically manifest as truncated reads that collapse into ISM transcript models. Because ISM classifications may also reflect genuine alternative TSS/TTS usage or incomplete sequencing [19], distinguishing biologically meaningful isoforms from degradation-associated fragments remains a major challenge during transcriptome curation [6]. Importantly, transcriptome reconstruction pipelines collapse multiple reads into representative transcript models, potentially masking systematic fragmentation patterns that are only apparent at the read-alignment level [30–32]. To investigate this problem, we used SQANTI3 [7] and SQANTI-reads [17] to analyze unprocessed reads from a direct-RNA Oxford Nanopore Technologies (ONT) degradation series spanning samples with progressively reduced RNA integrity number (RIN) [33].

Visual inspection in SQANTI-browser revealed a characteristic staircase-like pattern of progressively shortened 5′ transcript ends, consistent with degradation-driven fragmentation rather than regulated promoter usage (Fig. 3a). This organization was immediately apparent when visualizing reads within the browser, but was not readily identifiable from transcript classification tables alone. To distinguish genuine alternative TSS isoforms from degradation-associated fragments, we used the SQANTI-browser interactive filtering interface within the UCSC Genome Browser to subset transcripts whose TSS was located within +/-50 bp of experimentally supported FANTOM5 CAGE peaks (Fig. 3b). The threshold was selected empirically to illustrate interactive curation workflows and can be dynamically modified within the browser interface (Fig. 3b). This filtering selectively removed degradation-associated transcript fragments lacking promoter support while preserving reads supported by independent TSS evidence (Fig. 3c).

**Figure 3.**
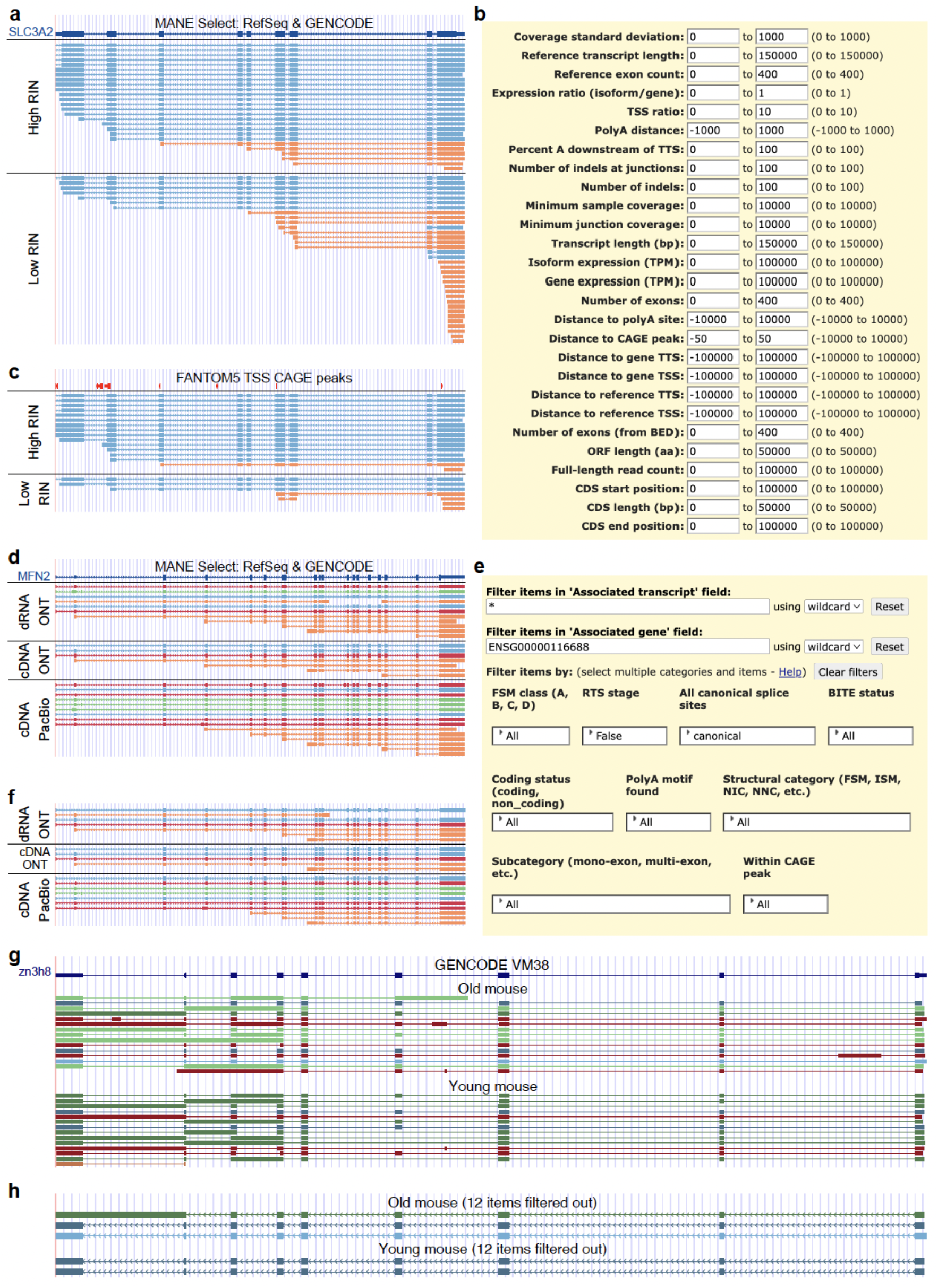
Interactive filtering of candidate isoforms in noisy loci. **a** Unfiltered direct RNA-sequencing showing incomplete splice match (ISM: orange) reads with staggered 5’ ends resulting from RNA degradation. **b** SQANTI-browser-defined filtering interface within the UCSC Genome Browser used to apply distance-based threshold relative to FANTOM5 CAGE peaks. **c** Curated view of the same locus where transcripts lacking transcription start site (TSS) functional support have been interactively removed. **d** Unfiltered comparison of direct RNA and cDNA Oxford Nanopore (ONT) and Pacific Biosciences (PacBio) datasets, including platform-specific inconsistencies. **e** SQANTI-browser-defined filtering interface within the UCSC Genome Browser used to apply categorical filters based on gene name, reverse transcriptase template switching (RTS), and canonical splice sites. **f** Refined ONT and PacBio transcriptomes after removing unsupported models. **g** Unfiltered view of the *Zc3h8* locus in old and young mice, exhibiting a high density of low-abundance novel in catalog (NIC: green) and novel not in catalog (NNC: red) transcripts. Dark colors indicate high transcript expression. **h** Filtered view of the same locus, where the old mouse has a high-confidence age-specific NIC isoform as well as full splice match (FSM: blue) isoforms, and the young mouse has FSM isoforms. Dark colors indicate high transcript expression.

Next, we evaluated SQANTI-browser’s capacity to resolve platform-specific sequencing biases. The Long-read RNA-Seq Genome Annotation Assessment Project (LRGASP) demonstrated that different sequencing chemistries introduce distinct profiles of transcriptomic “pseudo-novelty” [5]. ONT protocols, particularly direct-RNA sequencing, are more susceptible to premature termination, whereas cDNA-based protocols from both ONT and Pacific Biosciences (PacBio) are prone to RTS [5]. To isolate sequencing chemistry effects from assembly-driven artifacts, we analyzed the LRGASP consolidated transcriptome after restricting the dataset to transcript models independently reconstructed by at least three different algorithms.

Despite this algorithmic consensus, visual inspection of the *MFN2* locus still revealed substantial technical noise (Fig. 3d). Both ONT and PacBio datasets exhibited extensive 5’-truncated ISM models together with multiple low-abundance transcripts displaying irregular splice patterns. Therefore, we applied interactive SQANTI-browser filters to retain only transcripts initiating within +/-50 bp of validated FANTOM5 CAGE peaks, containing exclusively canonical splice junctions, and lacking evidence of RTS (Fig. 3e). These filtering criteria substantially reduced transcriptomic noise and produced a highly concordant set of core transcript models across all sequencing platforms (Fig. 3f). Notably, PacBio datasets retained a larger number of novel isoforms after filtering, likely reflecting the higher alignment precision of HiFi reads and the greater sensitivity of amplified cDNA libraries for low-abundance transcript detection relative to direct-RNA sequencing.

Finally, we evaluated the browser’s utility in distinguishing age-related transcriptomic variation from background noise. As splicing fidelity decreases during aging [34], transcriptomes from aged tissues frequently contain large numbers of low-abundance novel isoforms whose biological relevance is difficult to interpret. Thus, we analyzed the *Zc3h8* locus in long-read transcriptomes from young and old mouse brains [35]. In the unfiltered data, both samples had highly cluttered transcriptomes containing numerous low-abundance NIC and NNC transcript models (Fig. 3g).

To isolate high-confidence transcript structures, we applied stringent SQANTI-browser filters requiring TSS positions within +/-50 bp of FANTOM5 CAGE peaks, a minimum of two full-length supporting reads, exclusively canonical splice junctions, and absence of RTS or NMD annotations. These filters effectively cleaned the locus. In the young mouse, filtering removed all novel transcript models, leaving only annotated FSM isoforms. In contrast, the old mouse retained a high-confidence NIC isoform generated by an intron-retention event (Fig. 3h).

Taken together, these analyses demonstrate how SQANTI-browser enables interactive, evidence-guided curation of candidate isoforms across multiple sources of transcriptomic complexity. By integrating transcript structure, classification metadata, and orthogonal functional evidence within a unified visualization environment, SQANTI-browser facilitates the discrimination of technical artifacts from reproducible transcript diversity in large-scale long-read transcriptomes.

### Resolving splice-junction discrepancies and collaborative curation in non-reference genomes

A major challenge in long-read transcriptomics is the interpretation and dissemination of data generated from non-model organisms or newly assembled reference genomes that are absent from public repositories. SQANTI-browser addresses this limitation by dynamically generating UCSC-compatible Assembly Hubs that integrate non-reference genome sequences, transcript annotations, and SQANTI3-derived transcript classifications into a unified visualization environment. Because these hubs are hosted as standard web-accessible resources, complete browser sessions can be shared through persistent UCSC session links, allowing collaborators to access identical genomic coordinates, track configurations, and filtering states without requiring transfer of large alignment files or local browser configuration (Supplementary Fig. 2).

This framework also supports iterative transcriptome curation through integration between the SQANTI-browser interactive HTML reports and native UCSC utilities. Candidate isoforms can be filtered within the HTML interface according to structural category or quality metrics, and selected transcript identifiers can subsequently be exported through the “Generate Filter String” function. These identifiers can then be directly imported into the UCSC Table Browser to generate custom subtracks containing only the curated transcript subset (Supplementary Fig. 3). This workflow enables targeted inspection and sharing of specific transcript classes, including validated novel isoforms or disease-associated candidates, while preserving the broader transcriptome context.

To demonstrate these non-reference-genome capabilities while simultaneously resolving complex structural artifacts, we analyzed synthetic spike-in RNA variant (SIRV) controls [35, 36]. Because their exon–intron architecture is predefined, these transcripts provide a controlled benchmark in which structural discrepancies can be attributed to technical artifacts, alignment heuristics, or reference-related ambiguities rather than biological variation. Furthermore, as the SIRV reference genome is not available in the UCSC Genome Browser, this dataset illustrates SQANTI-browser’s ability to visualize transcriptomes derived from fully novel assemblies.

Inspection of reads and reconstructed transcript models derived from SIRV sequencing experiments using SQANTI-browser track visualization and interactive reports, revealed several transcripts classified as NNC despite originating from synthetic SIRV loci with well-defined reference structures (Fig. 4, Supplementary Fig. 4). Examination of these events within the SIRV4 locus revealed a systematic 4-bp splice-boundary shift at the SIRV403 and SIRV408 loci (Fig. 4a, Supplementary Fig 5a). This artifact was not apparent from tabular summaries alone, as their assigned subcategories included both “intron retention” and “at least one novel splice site” (Supplementary Fig 5b).

**Figure 4.**
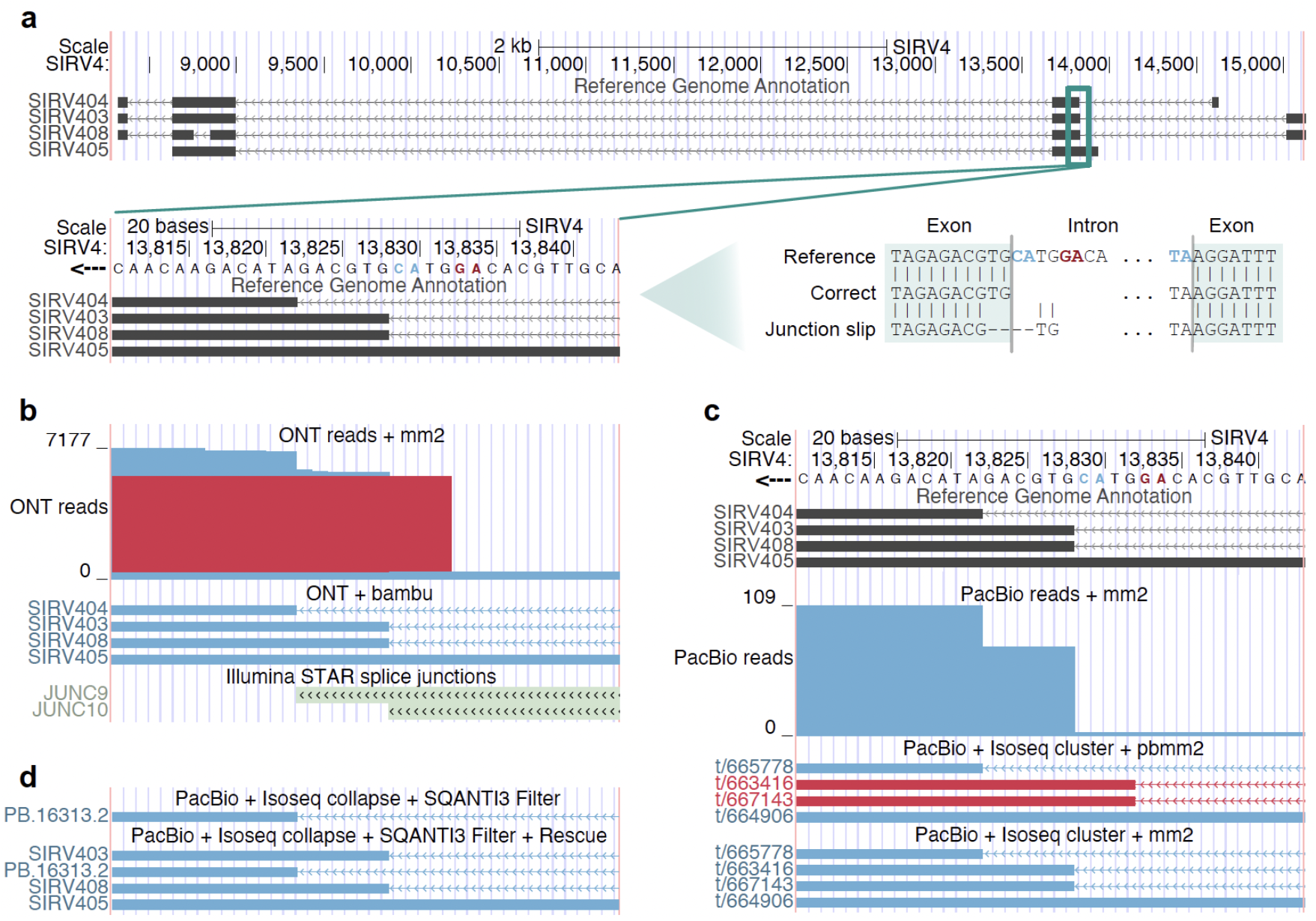
Splice junction discrepancies in synthetic SIRV transcriptomes. **a** Reference annotation of SIRV loci highlighting the complex splice junction. Inset: visualization of the acceptor splice motif, and sequence alignment showing both the reference junction and the alternative 4-bp shifted junction. Non-canonical splice motifs highlighted in red and canonical in blue. **b** Oxford Nanopore Technologies (ONT) read alignments at the locus classified as novel not in catalog (NNC: red), together with full splice match (FSM: blue) transcript models reconstructed with bambu and supporting Illumina short-read splice-junction evidence. **c** PacBio raw reads aligned with minimap2, and clustered isoforms mapped using pbmm2 and minimap2. FSM transcript models in blue and NNC in red. **d** Transcriptome curation illustrating the effects of the SQANTI3 machine-learning Filter and Rescue modules.

Inspection of the underlying genome sequence demonstrated that the Lexogen SIRV annotation defines the reference junction by an AT[A]..[T]AC splice signal. Although AT[A]..[T]AC junctions are canonical, they are extremely rare in mammalian transcriptomes (∼0.05%–0.06%) [37]. In contrast, alignment algorithms strongly favor highly prevalent GT[A/G]..[C/T]AG splice motifs and typically penalize rare or non-canonical transitions [38]. Within this sequence context, aligner heuristics favored a local 4-bp ‘junction slip’ that repositioned the acceptor site to generate a more favorable [T]AG boundary rather than preserving the annotated [T]AC acceptor (Fig. 4a).

Visualization using SQANTI-browser revealed that ONT reads aligned with minimap2 [38] were systematically extended by four nucleotides into the intronic space, resulting in the shifted splice junction classified by SQANTI3 as NNC (Fig. 4b, Supplementary Fig. 5-6). In contrast, when the same ONT reads were processed through the Bambu reconstruction pipeline [32], which incorporates splice-site modeling during transcript reconstruction, the resulting transcript models correctly recovered the annotated FSM structure. Independent Illumina short-read junction support generated with STAR [39] and integrated into the browser session further corroborated the reference-defined splice boundary (Fig. 4b).

We next evaluated whether the same ambiguity affected high-fidelity PacBio data. PacBio raw reads and de novo clustered isoforms generated independently of the reference [40] aligned correctly to the annotated splice site when mapped with the standard minimap2 [38] configuration (Fig. 4c). However, when clustered isoforms were aligned using pbmm2 within the official PacBio IsoSeq pipeline [40], the same 4-bp “junction slip” observed in the ONT data was systematically reproduced (Fig. 4c). Consequently, identical high-fidelity sequences were assigned fundamentally different structural categories (FSM vs. NNC) depending solely on the aligner-specific splice site heuristics. Nucleotide-level inspection within SQANTI-browser and the UCSC Genome Browser enabled direct tracing of this discrepancy to the alignment stage.

Finally, we evaluated the effect of automated filtering on these ambiguous transcript models. Application of the SQANTI3 machine-learning Filter [7] successfully removed the artifactual NNC isoforms generated by splice-boundary shifts, but also eliminated several high-confidence FSM transcripts, including SIRV405 (Fig. 4d). Therefore, we applied the SQANTI3 Rescue model, which incorporates orthogonal evidence to recover biologically supported isoforms removed during filtering. Visualization using SQANTI-browser confirmed that Rescue successfully restored the SIRV403, SIRV405, and SIRV408 FSM transcript models based on independent short-read splice-junction support (Fig. 4d).

Together, these analyses demonstrate how integrated visualization of transcript classifications, alignments, and orthogonal evidence via SQANTI-browser enables systematic resolution of alignment artifacts, reference ambiguities, and filtering discrepancies in non-reference transcriptomes.

### Scalability and computational performance of SQANTI-browser

To assess the computational efficiency and scalability of SQANTI-browser, we benchmarked its maximum memory consumption (RAM) and wall-clock execution time across transcriptomes of varying sizes, ranging from 10,000 to 1,000,000 subsampled isoforms, that had undergone SQANTI3 structural classification [7]. As illustrated in Supplementary Fig. 7, both memory usage and runtime had a linear relationship with the number of processed isoforms, demonstrating highly predictable and stable scaling. At the maximum tested scale of 1,000,000 isoforms, peak memory consumption was very low, at ∼1.1 GB. The pipeline finished in ∼75 seconds while restricted to a single CPU core. Because the maximum resource requirements (1.1 GB RAM, ∼75 seconds) fall well within the typical capabilities of standard personal computers, users can perform rapid, local curation of massive classification outputs with SQANTI-browser and the UCSC Genome Browser without relying on high-performance computing infrastructure.

## Discussion

The widespread adoption of long-read sequencing (LRS) has transformed transcriptomics from a discipline centered on gene-level quantification into one increasingly focused on isoform-resolved biology and transcript discovery [1]. As sequencing throughput and transcriptome reconstruction methods continue to improve, the major analytical challenge is no longer the generation of full-length transcript models, but the rigorous interpretation of structural novelty and the discrimination of biological signal from technical noise [6]. Existing reconstruction [30–32] and classification frameworks, including SQANTI3 [7], are indispensable for processing the scale and complexity of modern LRS datasets. However, these pipelines necessarily abstract millions of individual reads and probabilistic alignments into summarized transcript models and tabular classifications. Although computationally essential, this abstraction layer can separate transcript annotations from the underlying evidence required for expert interpretation.

SQANTI-browser was developed to bridge this gap by integrating transcript structural classifications, quality metrics, and orthogonal evidence directly within the UCSC Genome Browser environment [10]. Rather than treating transcript classification as a terminal computational output, SQANTI-browser enables iterative, evidence-guided curation in which transcript models can be evaluated in their genomic context alongside supporting experimental and population-scale datasets. Importantly, the framework preserves the full SQANTI3 [7] attribute space within indexed browser tracks, enabling dynamic filtering, attribute-aware querying, and interactive exploration without requiring external data manipulation or custom visualization infrastructure.

Genome browsers remain central to transcriptome curation because visual inspection provides a critical layer of validation that cannot be fully captured through automated classification alone. Expert curators routinely integrate orthogonal evidence—including splice-junction support, transcription start sites, conservation, and population-scale variation—to evaluate whether candidate isoforms are biologically plausible or technically derived. Existing genome browsers facilitate this process by aggregating diverse genomic datasets; however, they do not natively incorporate the extensive long-read-specific structural descriptors generated during transcriptome reconstruction. SQANTI-browser extends this evaluative framework by embedding the comprehensive SQANTI classification [7, 19] space directly into the visualization layer, allowing transcript novelty, structural complexity, and quality-control metrics to be interrogated simultaneously within their genomic context. As long-read sequencing expands into biodiversity genomics, clinical transcriptomics, and large-scale population studies, the ability to perform evidence-guided curation across custom transcriptomes and non-reference genomes will become increasingly important.

Our analyses demonstrate that this integrated visualization strategy is particularly valuable in loci affected by technical ambiguity or platform-specific artifacts. In the synthetic SIRV control transcriptome [36], the SQANTI-browser enabled nucleotide-level inspection of a systematic 4-bp splice-junction shift affecting the SIRV403 and SIRV408 loci. Although the underlying synthetic transcript structures were invariant, the resulting transcript classifications differed depending on the sequencing platform, alignment strategy, and reconstruction workflow used. The shifted splice junction was observed in ONT reads aligned with minimap2 [38] and in PacBio clustered isoforms aligned with pbmm2, whereas PacBio reads aligned with the standard minimap2 configuration and transcript models reconstructed with bambu [32] preferentially recovered the reference-defined structure. These discrepancies are consistent with heuristic alignment preferences favoring highly prevalent splice-site motifs during alignment scoring, illustrating how rare but valid splice boundaries can be misinterpreted as novel transcript structures. Critically, these artifacts were not readily identifiable from transcript classification tables alone and became apparent only through integrated visualization of raw read alignments, transcript models, splice-site sequence context, and orthogonal short-read evidence.

Beyond synthetic controls, our analyses of RNA degradation series [33], LRGASP benchmarking datasets [5], and aging mouse transcriptomes [35] further demonstrate that transcript novelty frequently emerges from a complex interplay between sequencing chemistry, library preparation biases, and biological heterogeneity. Direct-RNA ONT datasets in the degradation experiment exhibited pervasive 5′ fragmentation patterns that collapsed into ISM transcript models, whereas PacBio cDNA libraries also contained transcript models consistent with reverse transcriptase artifacts and amplification-associated complexity. Even after restricting analyses to transcript models reconstructed by multiple independent algorithms, substantial platform-specific “pseudo-novelty” remained detectable. These findings reinforce the limitation of relying exclusively on transcript-level consensus or automated filtering to define biological validity.

Importantly, our results also illustrate that aggressive filtering strategies can inadvertently remove biologically meaningful isoforms. In the *ZNF142* locus, classification-aware filtering combined with orthogonal evidence from GTEx splice junctions [25], SpliceAI predictions [26], evolutionary conservation [15], and population-scale sequencing data [16] enabled the identification of a high-confidence novel isoform absent from standard reference annotations. Similarly, in the aging mouse cortex, stringent evidence-guided filtering removed extensive transcriptomic noise while preserving a high-confidence age-associated intron-retention event in *Zc3h8*. Together, these examples demonstrate that biologically relevant transcript diversity may reside within structural classes that are routinely deprioritized during automated quality control. SQANTI-browser therefore shifts transcriptome curation from a purely exclusion-based paradigm toward an evidence-guided framework in which potentially meaningful isoforms can be systematically re-evaluated and rescued.

A central advantage of SQANTI-browser is that it leverages the existing UCSC Genome Browser ecosystem [10] rather than introducing a new standalone visualization platform. By producing fully compliant Track Hubs, the framework allows transcriptomes to be immediately contextualized alongside extensive public genomic resources, including conservation tracks, epigenomic annotations, population variation datasets, and predictive splicing models. This interoperability substantially lowers the barrier for transcriptome dissemination, collaborative review, and reproducible curation. Because SQANTI-browser outputs standard UCSC-style Track Hubs, transcriptomes can also be visualized in other compatible genome browsers such as Ensembl [12]. Although Ensembl currently supports only a subset of UCSC Track Hub functionality—including visualization of transcript structures and embedded metadata, but not dynamic filtering—this compatibility further expands interoperability across genomic platforms. In addition, because the framework operates directly on non-reference genomes and user-defined annotations, it remains applicable to synthetic transcriptomes, non-model organisms, and emerging experimental systems where standard genome-browser support may be limited.

We acknowledge several limitations of the current implementation. Although indexed bigBed tracks provide efficient visualization of large transcriptomes, browser-based rendering remains constrained when simultaneously displaying large population-scale cohorts or dense sequencing read datasets. Furthermore, because SQANTI-browser builds upon the UCSC Genome Browser infrastructure [10], visualization capabilities are inherently coupled to the constraints and feature set of the UCSC ecosystem. Nonetheless, the adaptive AutoSQL architecture was specifically designed to accommodate evolving metadata structures and emerging long-read applications. As single-cell and spatial long-read transcriptomics mature, the ability to integrate cell-type-specific, spatial, and multi-omic metadata within a unified visualization framework will become increasingly important.

Collectively, our results demonstrate that transcriptome interpretation cannot rely exclusively on automated classification outputs. Instead, robust curation of long-read transcriptomes requires integrated inspection of transcript structure, alignment behavior, orthogonal functional evidence, and genomic context. SQANTI-browser provides a scalable and reproducible framework for this process, transforming transcript classification from a static annotation task into an interactive, evidence-guided analytical workflow.

## Conclusion

SQANTI-browser enables interactive, classification-aware visualization and curation of long-read transcriptomes within the UCSC Genome Browser ecosystem. By embedding SQANTI3 structural classifications together with orthogonal validation data directly into indexed browser tracks, the framework allows researchers to systematically inspect transcript models, evaluate sequencing artifacts, and resolve structural ambiguity in genomic context. Through applications spanning clinical transcript discovery, noisy biological datasets, and synthetic benchmark controls, we demonstrate that integrated visualization substantially improves the interpretation of transcript novelty and transcriptome quality. SQANTI-browser therefore provides a scalable and interoperable framework for the transparent curation, dissemination, and benchmarking of modern isoform-resolved transcriptomes.

## Methods

SQANTI-browser is implemented as a Python command-line tool that converts SQANTI3 transcript classification outputs into fully configured UCSC Track Hubs. The software leverages the pandas library for scalable manipulation of large classification tables and uses Python’s subprocess interface to orchestrate UCSC command-line utilities for annotation conversion, indexing, and binary track generation. These utilities include tools for GTF-to-GenePred transformation, BED generation, BigBed compression, indexing, and 2-bit genome conversion.

The pipeline automates end-to-end Track Hub generation from SQANTI3 outputs, eliminating manual format conversion and configuration steps. To ensure reproducibility and portability, dependencies are managed through Conda, and containerized distributions are provided via Docker and Singularity.

All results presented in this manuscript were generated using default parameters in version 1.1.1 of SQANTI-browser. For a more detailed explanation and extensive documentation of the tool, please refer to our GitHub repository: https://github.com/ConesaLab/SQANTI-browser

### Transcriptome parsing and dynamic metadata

The primary inputs for SQANTI-browser’s main script (*sqanti_browser*.*py*) are the curated transcriptome (GTF) and the classification metadata (*classification*.*txt*) generated by SQANTI3 (quality control, filter, or rescue modules). Transcript coordinates are first extracted from the GTF file and converted into GenePred BED format using UCSC’s *gtfToGenePred* to standardize exon-intron structures. Transcripts are stratified into ten tracks based on SQANTI3 structural categories (FSM, ISM, NIC, NNC, Genic, Antisense, Fusion, Intergenic, and Genic_Intron) and a general track containing all transcripts. Category-specific color encoding is applied to facilitate rapid visual discrimination of transcript classes, and expression-aware shading highlights the highest expressed isoform per gene. BED files containing both positional data and color-coding are then merged with metadata from the classification file.

A core feature of SQANTI-browser is its adaptive metadata modeling. Instead of using a hardcoded schema, the tool inspects the input classification file to identify all available metadata fields. It then programmatically generates a custom AutoSQL definition extending the BED12 specification, producing BED12+N BigBed files, where N corresponds to the number of metadata columns in the SQANTI3 output (e.g., 48 fields in SQANTI3 v6.0.0). This dynamic schema generation ensures forward compatibility with evolving versions of the SQANTI3 pipeline without requiring internal modifications to the software’s core visualization framework. Furthermore, this adaptive attribute mapping allows for the seamless inclusion of non-standard descriptors—such as single-cell cluster identifiers or spatial coordinates—provided they are present in the input classification file, thereby future-proofing the tool for emerging multi-omic and high-resolution transcriptomic applications.

The resulting BED files are sorted following the user-defined requirement by the *--sort-by* flag, and to ensure scalability for large transcriptomes, BED files are compressed into binary indexed BigBed (.bb) format using UCSC’s *bedToBigBed*.

### Integration of orthogonal validation data and non-reference genomes

To support evidence-driven curation, SQANTI-browser integrates orthogonal data as independent browser tracks. These include splice junction (SJ.out.tab) evidence derived from short-read mapping using STAR, TSS information from CAGE-seq, and PolyA sites (TTS) annotations from Quant-seq assays. Input files in BED or tabular formats are converted into indexed BigBed tracks for consistent visualization and coordinate-based integration.

User-supplied reference annotations (GTF) are likewise converted into distinct BigBed tracks to facilitate locus-specific comparison between reconstructed transcript models and reference gene structures. This layered configuration enables direct assessment of splice junction concordance, transcript boundary support, and structural novelty within a unified genomic coordinate system.

For non-model organisms, in-house assemblies, or synthetic sequences (e.g., SIRVs [36] or Sequins [41]), SQANTI-browser automatically generates fully configured UCSC Assembly Hubs. When provided with FASTA assemblies, the tool orchestrates the conversion to 2-bit format using UCSC’s *faToTwoBit* and compiles the necessary genomes.txt and trackDb.txt configuration files to define the custom genomic coordinate system. This functionality enables the visualization of transcriptomes on genomes not natively hosted by the UCSC Genome Browser, ensuring that the tool’s classification-aware filtering and search capabilities remain fully operational on synthetic or alternative assemblies.

### Metadata-aware filtering, free-text querying, and interactive reporting

SQANTI-browser embeds the complete SQANTI3 classification matrix directly within the visualization layer. Filter definitions are automatically generated within the Track Hub configuration, in the trackDb.txt configuration file, enabling server-side filtering of categorical variables (e.g., *filterValues*.*structural_category*) and numerical attributes (e.g., *filterByRange*.*FL, filterLimits*.*iso_exp*) through the UCSC interface. Users can dynamically subset transcript models based on structural category, expression metrics, junction support, coding potential, or additional quality descriptors encoded in the metadata.

To support free-text querying, SQANTI-browser constructs a Trix-based searchable index from the classification matrix. Indexed metadata allow keyword-based retrieval of isoform subsets directly within the UCSC Genome Browser search bar (e.g., “GeneA NNC coding strand_+”), enabling rapid identification of transcripts matching user-defined combinations of attributes.

Optionally, when the *--tables* flag is enabled, SQANTI-browser generates standalone interactive HTML5/JavaScript-based reports summarizing transcriptome metadata. These reports provide local, searchable, and exportable tables, facilitating offline curation and reproducible selection of isoform subsets. Furthermore, the interactive reports provide a ‘Generate Trix String’ button to produce ready-to-paste query strings in the Trix Keyword search bar, to reproduce the search within the UCSC Genome Browser, and users can select rows of interest in these tables and use the ‘Generate Filter String’ button to obtain a newline-separated list of isoform identifiers to extract a custom track containing only the selected isoforms, enabling locus-specific inspection of curated subsets without modifying the full hub.

### Track Hub configuration and deployment

The final output is compiled into a portable UCSC Track Hub directory conforming to official UCSC specifications. The output includes indexed BigBed files, hub configuration files (Hub.txt and trackDb.txt), and superTrack containers to group all SQANTI-derived tracks. This Track Hub configuration defines visibility priorities, filter definitions, search index associations, and HTML description pages. The resulting hub can be deployed on any static web-accessible server (e.g. GitHub, institutional web space, or cloud storage with public read access) and loaded into the UCSC Genome Browser without additional server-side configuration.

For multi-sample experiments (e.g. when processing SQANTI-reads outputs [17]), the optional *--hub-name* parameter allows sample-specific track labeling and namespace prefixing (e.g. ‘Sample1 SQANTI3 Transcripts’), enabling simultaneous loading of multiple hubs within a single UCSC session. This supports direct locus-by-locus comparison of transcript structure and classification across samples without naming conflicts.

## Visualization of real data using SQANTI-browser

To demonstrate the utility of SQANTI-browser for transcript filtering, exploration, and visualization, we applied it to five representative long-read transcriptomics datasets (i) the isoform-centric microglia genomic atlas [18], (ii) direct-RNA ONT sequencing from an RNA degradation experiment [33], (iii) ONT and PacBio sequencing data from the human induced pluripotent stem cell line WTC11 generated by the LRGASP consortium [5], (iv) transcriptomes from young and old mice [35], and (v) sequencing data from SIRV control mixes produced by Lexogen [35, 36].

### Visualization of clinical transcripts from the isoform-centric microglia genomic atlas

Human microglial long-read transcriptomic data were obtained from the isoform-centric microglia genomic atlas [18]. We utilized the high-quality, processed GTF file provided by the study, which includes transcript models derived from 30 post-mortem human brain samples. These data were originally generated using PacBio Sequel II long-read sequencing to elucidate disease-associated genetic regulation of splicing in microglial cells.

Detailed data processing information is available in [18]. Briefly, the standard PacBio processing pipeline was used, including PacBio CCS (v4.0) raw file collapsing, LIMA (v1.11) barcode removal, isoseq3 refine (v3.2) trimming of poly(A) and poly(T) tails, and pbmm2 (v1.4.0) (https://isoseq.how/) mapping to the hg38 reference genome. Transcript reconstruction was performed using StringTie2 (v2.2.1) [31] in hybrid assembly mode, combining long- and short-read samples using the *--mix* argument.

SQANTI3 (v6.0.1) [7] was applied to characterize the provided transcriptome, incorporating ORF prediction and coding potential evaluation using the --include_ORF argument. Using the *--tables* argument, SQANTI-browser produced interactive tables in HTML5 format that were used to identify transcripts of interest by filtering for predicted coding potential by TransDecoder2 and PSAURON and high positional concordance (+/-50 bp) with reference FANTOM5 CAGE peaks (TSS) [13] and PolyASite Atlas (TTS) [22] entries, while excluding transcripts flagged for RTS or non-canonical splice sites. For isoform curation, the generated hub was visualized in the UCSC Genome Browser together with native tracks from GENCODE v49 [2], FANTOM5 [13], recount3 GTEx [24, 25], SpliceAI [26], AbSplice [27], OMIM [23], gnomAD [16], RepeatMasker [28], 100 vertebrates conservation by PhyloP, and Multiz alignments of 100 vertebrates [10, 15].

### Visualization of RNA degradation in direct RNA ONT sequenced data

Direct-RNA sequencing data was obtained from a publicly available controlled RNA degradation experiment [33]. Total RNA was extracted from the K562 human cell line and subjected to sequential degradation at room temperature for 0, 4, 8, and 24 hours prior to sequencing using MinION and PromethION ONT sequencing. Raw reads obtained from the European Nucleotide Archive (ENA) were aligned to the hg38 reference genome using minimap2 (v2.28) [38] and *-ax splice -uf --MD -t 4* parameters [38]. Mapped transcriptomes were classified using SQANTI3 (v5.3.6) [7] and SQANTI-reads (v1.0) [17]. Highly degraded loci were identified by calculating the ratio of ISM reads between the samples with the highest and lowest RNA integrity numbers (9.8 and 7.7, respectively). Degraded isoforms were curated within the UCSC Genome Browser by defining proximity of +/-50 bp to experimentally validated FANTOM5 CAGE peaks.

### Visualization of platform-specific artifacts from ONT and PacBio data

SQANTI-browser was applied to long-read transcriptome data from the WTC11 human cell line generated within the LRGASP consortium [5]. We utilized the high-quality, processed BED file provided by the study, which includes consolidated transcriptomes resulting from 6 combinations of library preparation strategies, 2 sequencing platforms, and 13 different transcript reconstruction tools [5]. We subset 3 different GTF files from the consolidated BED, each representing cDNA-PacBio, cDNA-ONT, and dRNA-ONT combinations. After structural classification using SQANTI3 (v5.3.6) [7], we evaluated the technical discrepancies between the three strategies using SQANTI-browser. Candidate isoforms were interactively curated within the UCSC Genome Browser using the following parameters: proximity of +/-50 bp to experimentally validated FANTOM5 CAGE peaks, all canonical splice junctions, and absence of the RTS flag.

### Visualization of age-associated novel isoforms

PacBio Sequel II sequenced data from the brains of three young (3-month-old) and two old (15-month-old) C57BL6/J male mice were obtained from [35]. Raw FASTQ files were aligned to the reference mm39 NCBI RefSeq genome using minimap2 (v2.28) [38] with *-ax splice:hq -uf* parameters. Transcriptome reconstruction was performed using TAMA Collapse [30] with options *-a 50 -m 0 -z 50* to ensure that only reads with the same splice junctions were collapsed into the same transcript model, while leaving a threshold of 50 bp for TTS and TSS to account for transcript end degradation. Next, SQANTI3 (v5.3.6) was used for both structural characterization and quality filtering via the rules module [5]. The filtering criteria were as follows: FSM transcripts were retained unless a stretch of nucleotides with at least 60% adenines was detected after the TTS, which would indicate an unreliable 3’ end due to potential intrapriming. Other transcript categories were retained if no intrapriming was detected, no junction was labelled as RTS, and all junctions had a minimum coverage of one read by Illumina short reads. Age-related noisy loci were identified by calculating the ratio of NIC and NNC isoforms between the old and young mice samples.

The resulting filtered transcriptomes were evaluated using SQANTI-browser. Candidate isoforms were interactively curated within the UCSC Genome Browser using the following parameters: proximity of +/-50 bp to experimentally validated FANTOM5 CAGE peaks, a minimum of two full-length read counts, all canonical splice junctions, and the absence of RTS and NMD flags.

### Visualization of the SIRV genome

To demonstrate the ability of SQANTI-browser to operate on non-reference genomes, non-model organisms, and annotations, we generated a UCSC Track Hub for the SIRV E0 control set (Lexogen) [36].

LRS datasets were obtained from the publicly available mouse brain experiment containing SIRV spike-ins [35]. Raw ONT and PacBio HiFi reads were aligned to the SIRV reference genome using minimap2 (v2.28) [38] with *-ax splice -uf* and *-ax splice:hq -uf* parameters, respectively. For PacBio data, high-fidelity reads were additionally processed using the IsoSeq3 pipeline (v4.3.0) with the *cluster* command to generate de novo isoform clusters. These clustered isoforms were subsequently aligned to the reference using both minimap2 (v2.28) [38] and pbmm2 (v1.17.0) (https://isoseq.how/) to evaluate potential discrepancies introduced by mapper-specific alignment.

Full-length transcript isoforms were reconstructed using bambu (v3.8.3) [32] for both ONT and PacBio dataset, and the IsoSeq3 pipeline for PacBio-specific analyses. The resulting transcript models were processed using SQANTI3 (v5.3.6) [7] to obtain structural classifications and quality-control metrics.

Quality control was performed through a two-stage procedure: 1) Automated filtering using the SQANTI3 machine-learning (ML) module to remove potential artifacts. 2) Evidence-guided recovery of high-confidence isoforms that were removed during ML filtering but retained strong expression or structural support were recovered using the SQANTI3 Rescue module [7].

All SQANTI3 classifications and associated metadata were integrated into a UCSC Track Hub using SQANTI-browser. Because the SIRV reference genome is not available in the UCSC Genome Browser, a non-reference assembly hub was generated using the --*twobit* option to automatically create the required reference sequence. Orthogonal validation evidence was incorporated using the *--star-sj* option in SQANTI-browser, which integrates splice junction support derived from Illumina short-read data. Short reads from the corresponding SIRV-spiked libraries were aligned using STAR (v2.7.11), and junction support was provided through the resulting SJ.out.tab files.

All alignment and transcript reconstruction analyses were performed using default parameters unless otherwise specified. Orthogonal visualizations of BAM files in the Integrative Genomics Viewer (v2.19.7) were generated to corroborate SQANTI-browser findings.

## Computational efficiency evaluation

To evaluate the scalability of SQANTI-browser, we generated five transcriptome datasets of decreasing size by randomly subsampling isoforms (1,000,000; 500,000; 100,000; 50,000; and 10,000 isoforms) from the SAMEA117622252 PacBio-sequenced sample available in the ENA, processed following the IsoSeq pipeline recommended by PacBio (https://isoseq.how/). Each subset was independently processed with SQANTI3 to generate the corresponding transcript annotations and classification metadata required as input for SQANTI-browser. This approach produced transcriptomes spanning a range of reconstructed isoform counts and classification file sizes.

Each dataset was processed using SQANTI-browser from classification input to complete UCSC Track Hub generation. All benchmarking experiments were executed on a Linux high-performance computing cluster, with jobs allocated a single CPU core and 4 GB of RAM.

Runtime was measured as wall-clock execution time from invocation to completion of the SQANTI-browser pipeline. Memory consumption was quantified using the Slurm *sacct* monitoring utility and */usr/bin/time -v*. These metrics were recorded for each dataset to characterize runtime and memory scaling as a function of transcriptome size.

## Abbreviations

TSS: transcription start site
TTS: transcription termination site
RTS: reverse transcriptase template switching
NNC: Novel not in catalog
ISM: incomplete splice match
FSM: full splice match
NIC: novel in catalog
NMD: nonsense-mediated decay
ONT: Oxford Nanopore Technologies
PacBio: Pacific Biosciences
SIRV: synthetic spike-in RNA variant
RAM: random access memory
RIN: RNA integrity number
ENA: european nucleotide archive
ML: machine learning

## Declaration

### Ethics approval and consent to participate

Not applicable.

### Consent for publication

Not applicable.

### Availability of data and materials

SQANTI-browser is findable at https://github.com/ConesaLab/SQANTI-browser and accessible via open-source GNU GPLv3 license. It is platform independent and mainly written in python and HTML. The SQANTI-browser source code used in this study is also accessible as a software release: https://github.com/ConesaLab/SQANTI-browser/releases/tag/v1.1.1. The human reference genome sequence (GRCh38) and gene annotation v43 from GENCODE can be obtained from https://ftp.ebi.ac.uk/pub/databases/gencode/Gencode_human/release_43. The direct RNA ONT sequencing data from the controlled RNA degradation experiment was obtained from the European Nucleotide Archive (ENA; https://www.ebi.ac.uk/ena/browser/home) under accession number PRJEB53210. The processed long-read sequencing data obtained from the human WTC11 cell line is available at https://cgl.gi.ucsc.edu/data/LRGASP/. The reference SIRV annotation was obtained from Lexogen (https://www.lexogen.com/sirvs/) and the reads of the E0 set for ONT, PacBio and Illumina from ENA under accession number PRJEB85167, where the young and old mouse brain data were available. The visualizations generated and analyzed during the current study can be reproduced using the code and example data provided in the https://github.com/carolinamonzo/figures_SQANTI-browser repository. Example SQANTI-browser-UCSC session with *ZNF142* NNC event in isoform of interest: https://genome.ucsc.edu/s/cmonzo/test_SQANTI%2Dbrowser

### Authors’ contributions

A.P., C.M., A.C. and M.D. conceptualized SQANTI-browser. C.M. developed and implemented the software. A.P., C.B., A.C.F. and C.M. performed the analysis and generated visualizations. A.P. C.B. and C.M. wrote the manuscript. C.M. designed the approach and supervised the work. All authors read and approved the final manuscript.

## Acknowledgements

The authors thank Pablo Atienza López for his artistic design of the SQANTI-browser logo, and Xanthi-Lida Katopodi for scientific discussion. We thank Isabel Fariñas and José Manuel Morante from the University of Valencia, and Luis Ferrández for mouse handling and data generation. The development of the tool and computations were performed on the high performance computing cluster Garnatxa at the Institute for Integrative Systems Biology (I^2^SysBio), I^2^SysBio is a mixed research centre formed by University of Valencia (UV) and Spanish National Research Council (CSIC).

