## Supplementary file for "SQANTI-browser: visualization and curation of SQANTI3-classified long-read transcriptomes within the UCSC Genome Browser"

SQANTI3 Analysis: novel\_not\_in\_catalog Isoforms


### SQANTI3 Analysis: novel\_not\_in\_catalog Isoforms

ReferenceNew SiteIsoform (NNC)

**Category Definition:** **NNC (Novel Not in Catalog):** The query isoform does not have a FSM or ISM match, and has at least one donor or acceptor site that is not annotated.

For more details on categories, visit here: SQANTI3 Wiki: Categories.
  
For a detailed explanation of each column of the table (SQANTI3 classification), see here: Column Glossary.

---

**How to Filter:**

- **Numeric Columns (e.g., length, exons):** Use ranges like `100:1000` (values between 100 and 1000), `100:` (>100), or `:1000` (<1000). Single numbers perform an exact match.
- **Dropdowns:** Select a value to filter for an exact match.
- **Text Columns:** Type to search for substrings (case-insensitive).

---

**Generate Trix String (for UCSC Genome Browser search):**

- Use dropdown filters and click "Generate Trix String" to get a search based on your filter criteria.
- *Note:* Range filters (e.g., `100:1000`) are not supported by Trix and will be ignored, only exact numeric matches are supported.
- Search terms automatically use underscores (e.g., `structural_category_full_splice_match`, `strand_plus`, `coding_coding`).
- Click rows to select (click again to deselect). Multiple rows can be selected. Then click "Generate Trix String" to get search terms.

**Generate Filter String (for Table Browser):**

- Select one or more rows (or apply filters to narrow the table), then click "Generate Filter String" to get a newline-separated list of isoform IDs.
- Copy the list and paste (one ID per line) into UCSC Table Browser when filtering the track by the `name` field to create a custom track with only those isoforms.

Total transcripts: 1838 | Close Window

| isoform | chrom | start | end | strand | length | exons | structural\_category | subcategory | FSM\_class | associated\_gene | associated\_transcript | ref\_length | ref\_exons | diff\_to\_TSS | diff\_to\_TTS | diff\_to\_TSS\_genomic | diff\_to\_TTS\_genomic | diff\_to\_gene\_TSS | diff\_to\_gene\_TTS | RTS\_stage | all\_canonical | min\_sample\_cov | min\_cov | min\_cov\_pos | sd\_cov | FL | n\_indels | n\_indels\_junc | bite | iso\_exp | gene\_exp | ratio\_exp | coding | ORF\_length | CDS\_length | protein\_length | CDS\_start | CDS\_end | CDS\_genomic\_start | CDS\_genomic\_end | psauron\_score | CDS\_type | predicted\_NMD | perc\_A\_downstream\_TTS | seq\_A\_downstream\_TTS | dist\_to\_CAGE\_peak | within\_CAGE\_peak | dist\_to\_polyA\_site | within\_polyA\_site | polyA\_motif | polyA\_dist | polyA\_motif\_found | ratio\_TSS |
| --- | --- | --- | --- | --- | --- | --- | --- | --- | --- | --- | --- | --- | --- | --- | --- | --- | --- | --- | --- | --- | --- | --- | --- | --- | --- | --- | --- | --- | --- | --- | --- | --- | --- | --- | --- | --- | --- | --- | --- | --- | --- | --- | --- | --- | --- | --- | --- | --- | --- | --- | --- | --- | --- |
| isoform | chrom | start | end | strand | length | exons | structural\_category | subcategory | FSM\_class | associated\_gene | associated\_transcript | ref\_length | ref\_exons | diff\_to\_TSS | diff\_to\_TTS | diff\_to\_TSS\_genomic | diff\_to\_TTS\_genomic | diff\_to\_gene\_TSS | diff\_to\_gene\_TTS | RTS\_stage | all\_canonical | min\_sample\_cov | min\_cov | min\_cov\_pos | sd\_cov | FL | n\_indels | n\_indels\_junc | bite | iso\_exp | gene\_exp | ratio\_exp | coding | ORF\_length | CDS\_length | protein\_length | CDS\_start | CDS\_end | CDS\_genomic\_start | CDS\_genomic\_end | psauron\_score | CDS\_type | predicted\_NMD | perc\_A\_downstream\_TTS | seq\_A\_downstream\_TTS | dist\_to\_CAGE\_peak | within\_CAGE\_peak | dist\_to\_polyA\_site | within\_polyA\_site | polyA\_motif | polyA\_dist | polyA\_motif\_found | ratio\_TSS |
