## Supplementary figures for "SQANTI-browser: visualization and curation of SQANTI3-classified long-read transcriptomes within the UCSC Genome Browser"

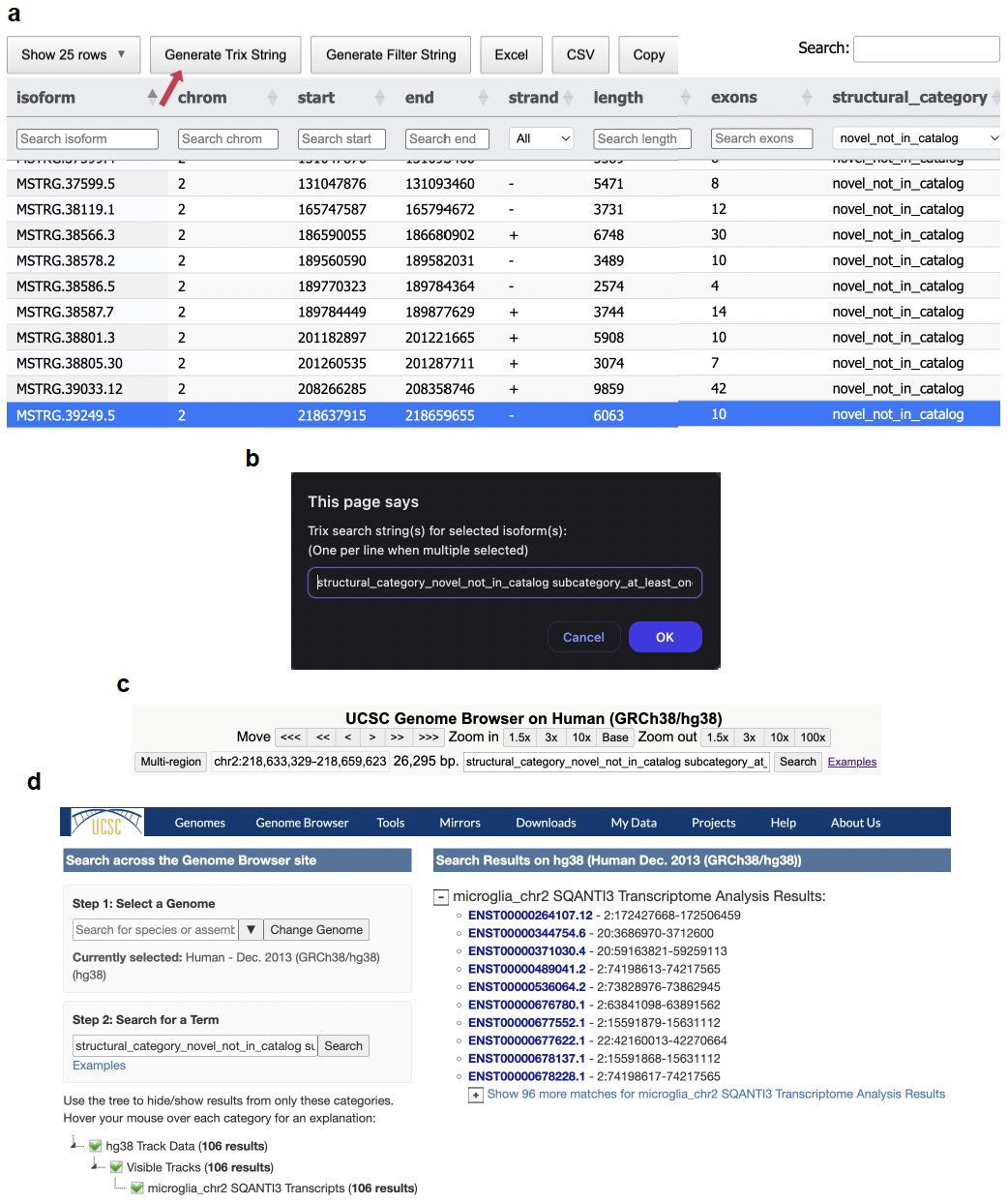

**Supplementary figure 1:** Visualization of Trix functionality. **a** SQANTI-browser’s HTML report showcasing the Generate Trix String button as well as selected filters and highlighted isoform of interest. **b** Trix string generated by “Generate Trix String” button. **c** Search of Trix string in the UCSC Genome Browser search bar. **d** Identified isoforms that match the criteria defined by the Trix string.

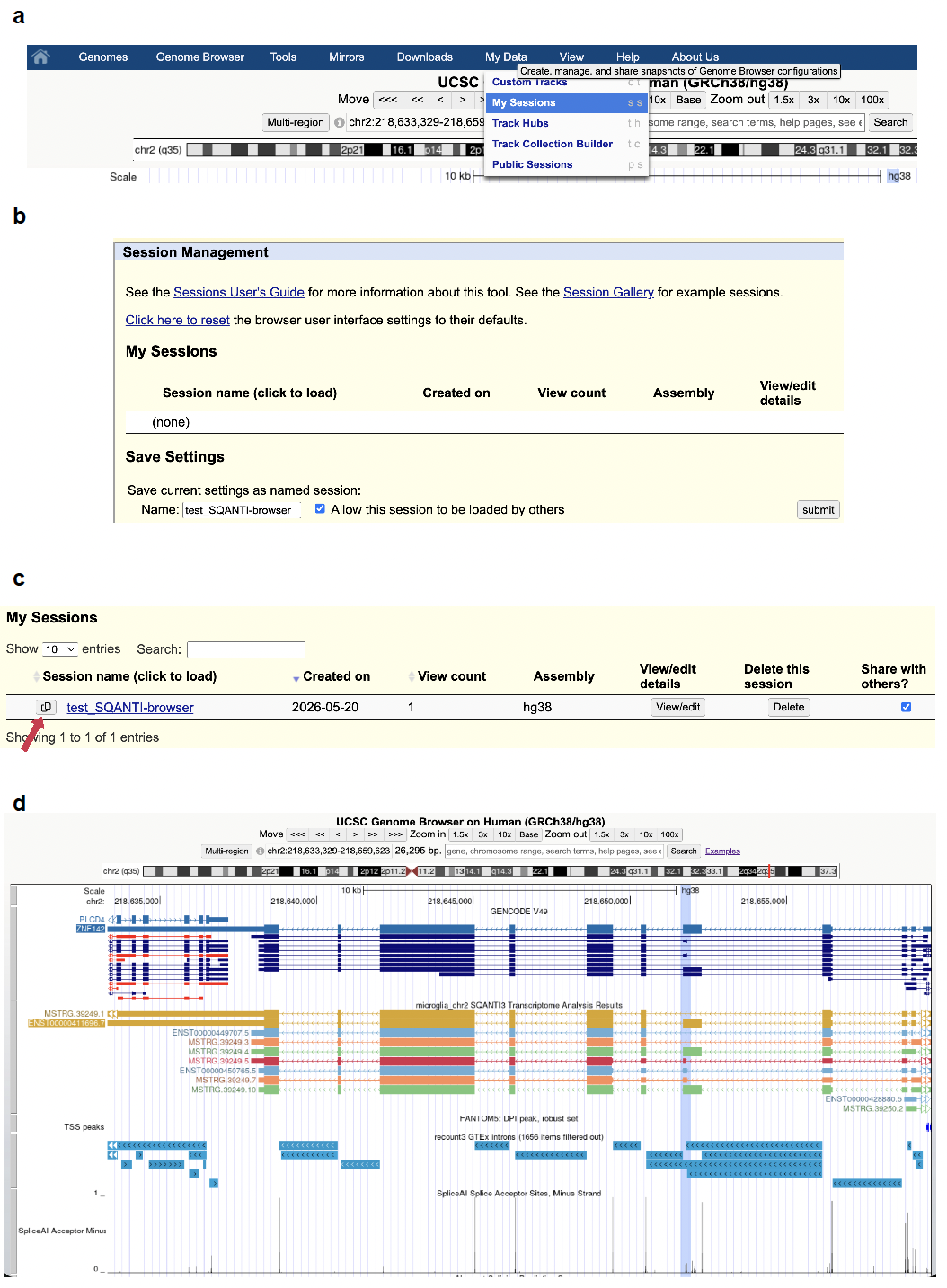

**Supplementary figure 2:** Generation of session links to share assembly hubs. **a** Selection of My Sessions in the UCSC Genome Browser. **b** My sessions functionality allows saving the current browser view, data and configuration into a shareable session. **c** A link to share session is generated and available to copy and share with collaborators. **d** Visualization of the shared session by opening the shared link in a navigator allows for identification of a highlighted region of interest for collaborative curation.

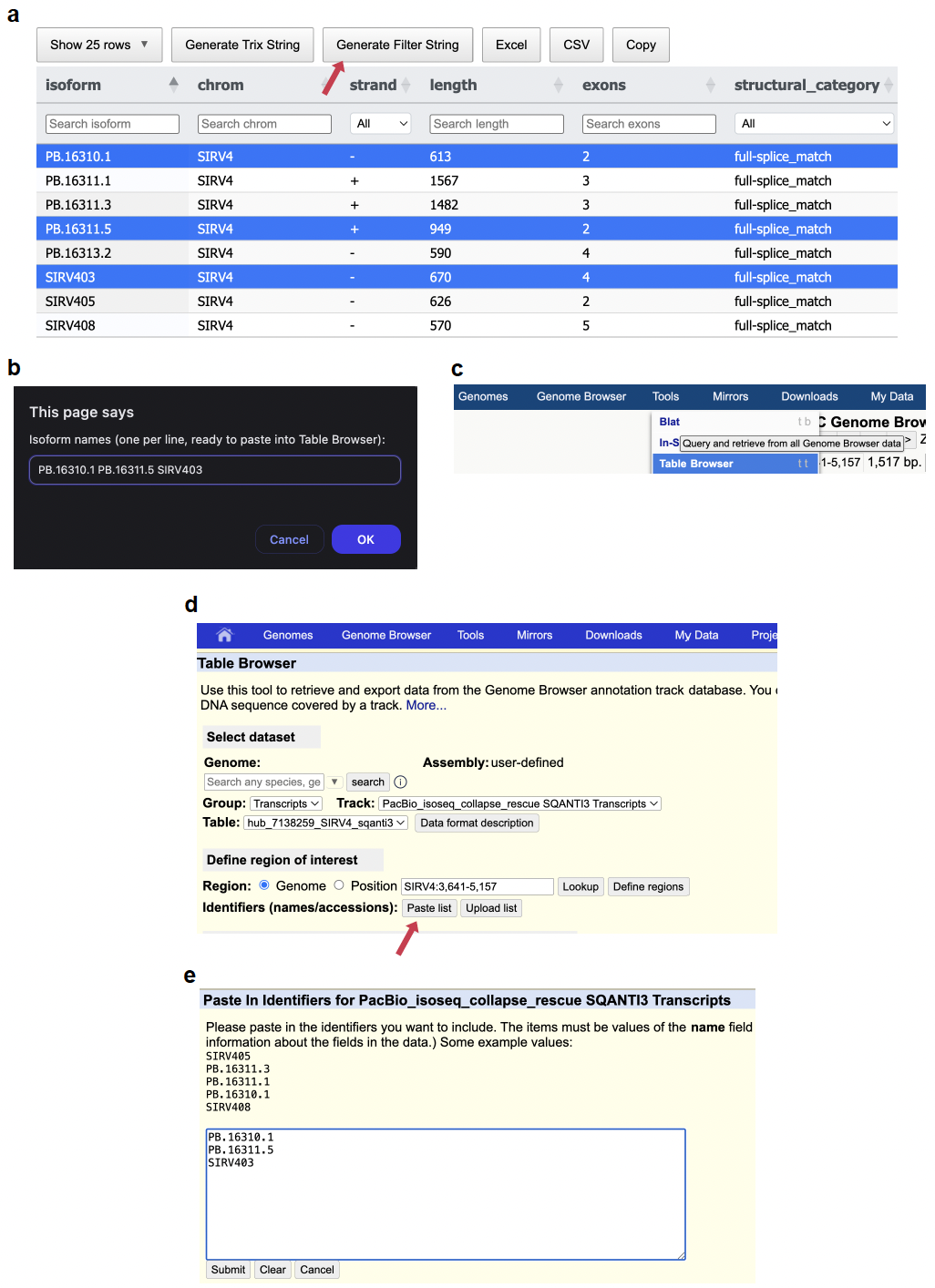

**Supplementary figure 3:** Visualization of Search String functionality. **a** Selected isoforms of interest in the SQANTI-browser HTML report. **b** String of isoform names of interest generated by “Generate Filter String” button. **c** Selection of Table Browser tool in the UCSC Genome Browser. **d** Table Browser functionality permits pasting a list of identifiers to subset. **e** Example pasted list of isoforms, ready to submit.

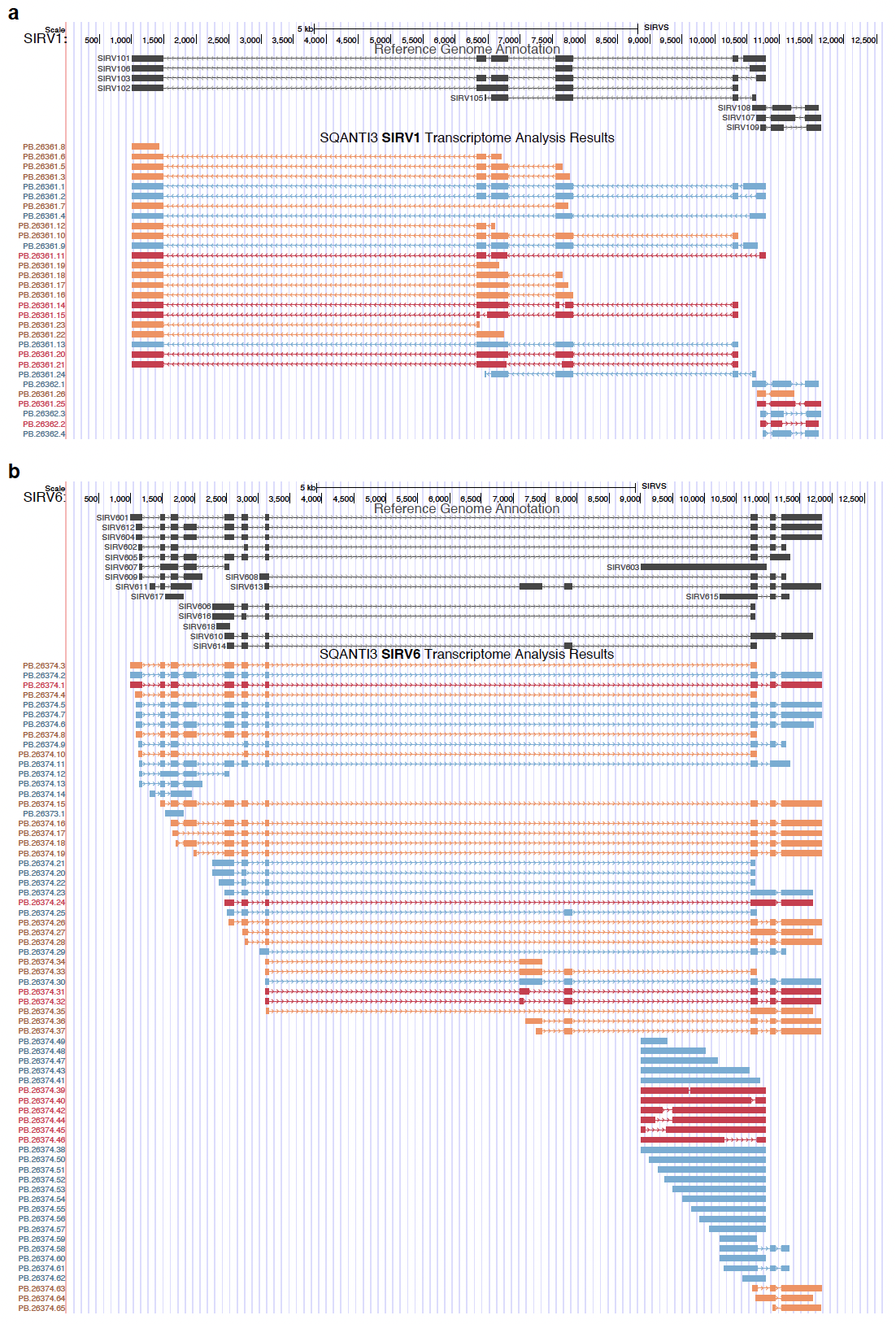

**Supplementary Figure 4:** Example visualization of **a** SIRV1 and **b** SIRV6 using SQANTI-browser. Red isoforms depict high presence of Novel Not in Catalog (NNC) isoforms in the synthetic SIRV molecules.

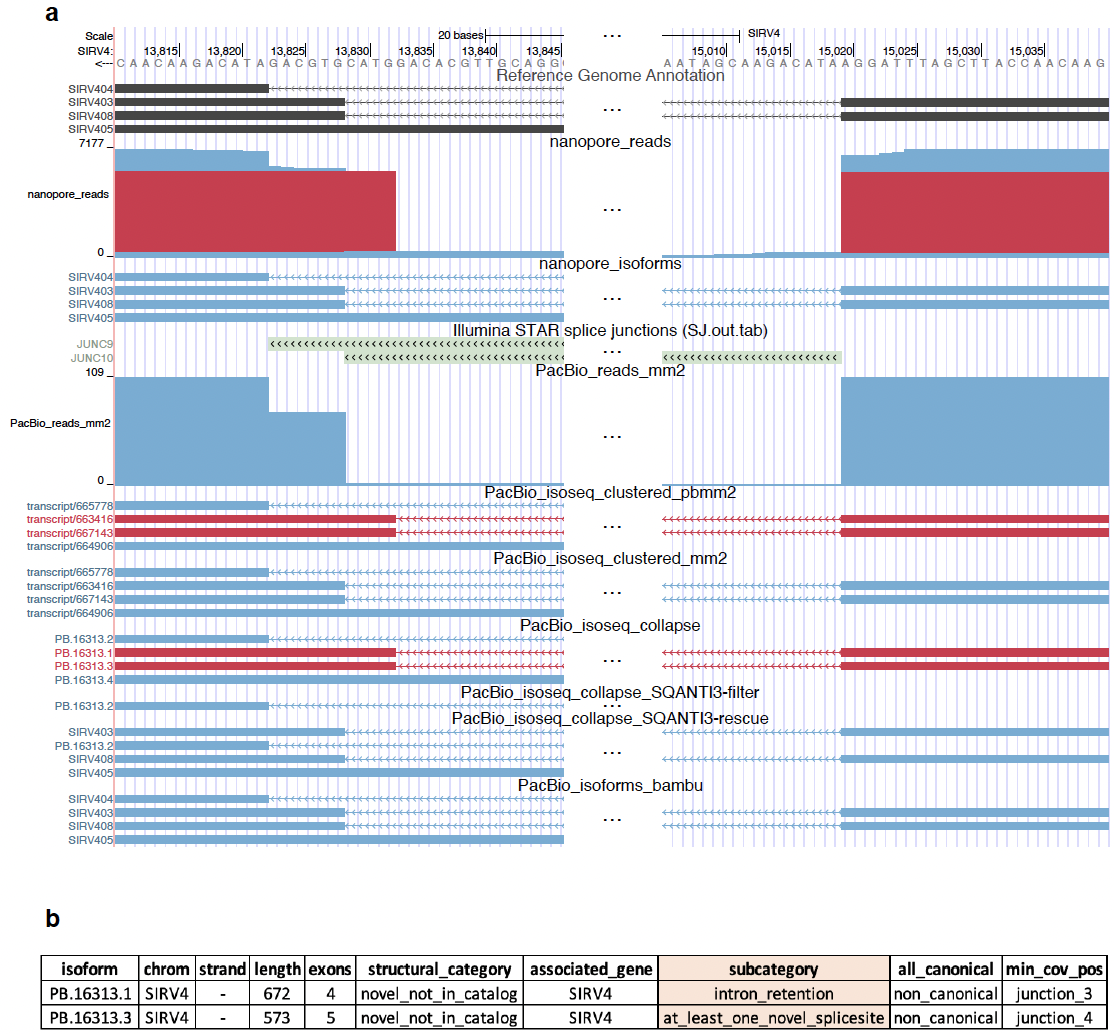

**Supplementary Figure 5:** Visualization of splice junction information in SIRV4. **a** SQANTI-browser view of the acceptor and donor sites, including all variations of the processing pipeline. **b** Exported csv from SQANTI-browser’s interactive HTML reports highlighting the different subcategories of the isoforms and reads in SIRV403 and SIRV408.

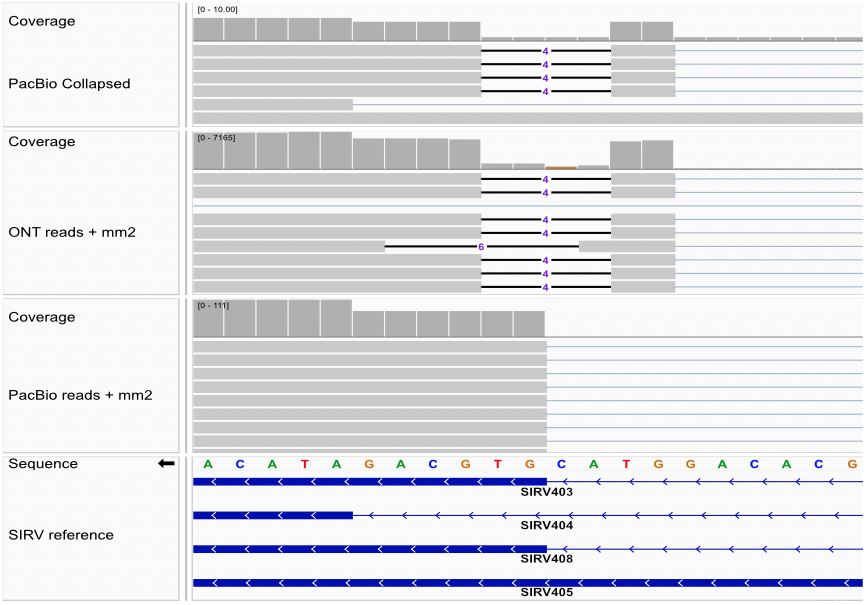

**Supplementary Figure 6:** Supporting visualization of the alternative 4-bp shifted junction using the Integrative Genomics Viewer on Oxford Nanopore Technologies (ONT) and Pacific Biosciences (PacBio) reads mapped with minimap2, and PacBio isoforms obtained via IsoSeq Collapse.

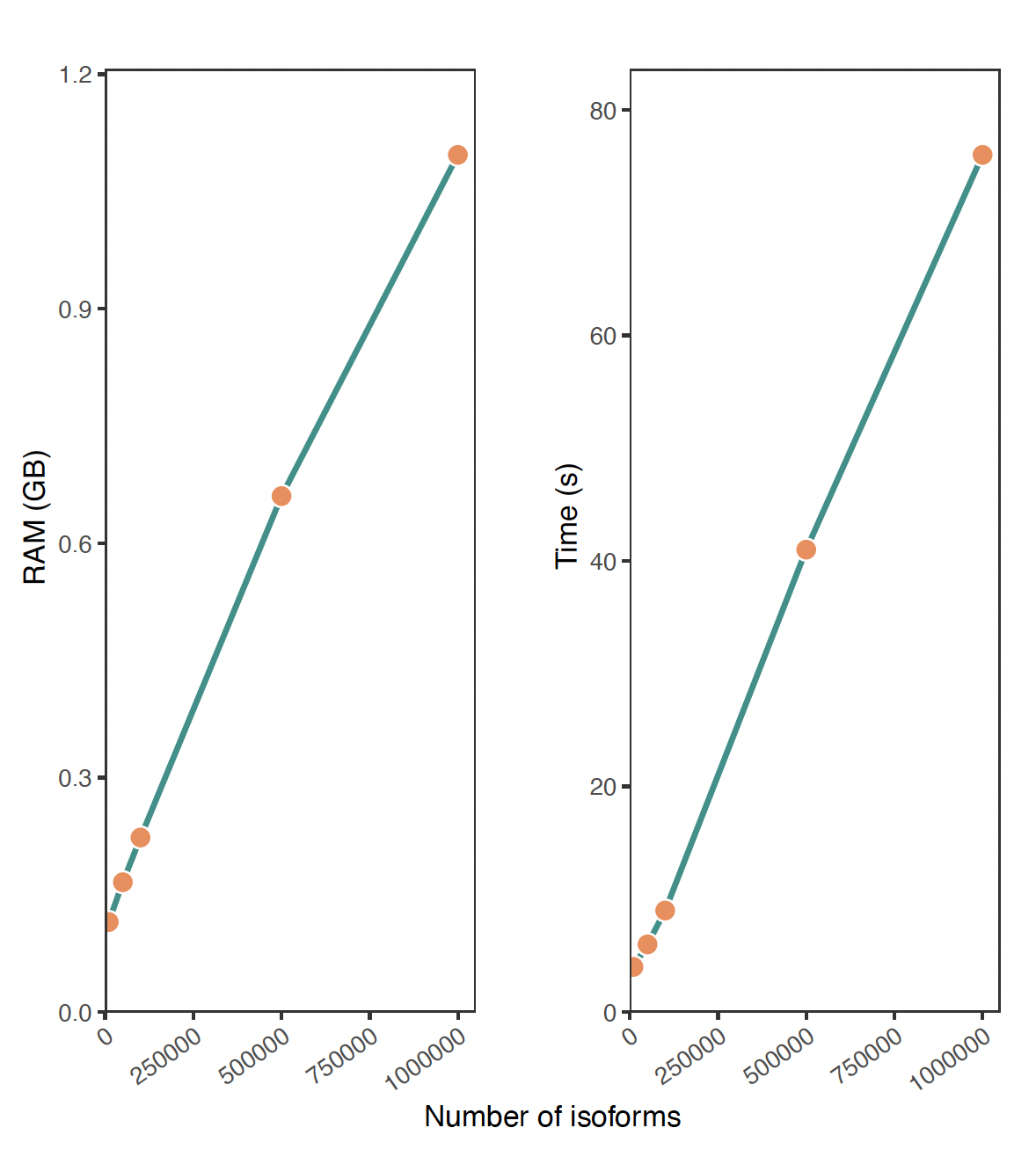

**Supplementary Figure 7:** Memory usage and runtime of SQANTI-browser ran on increasing amounts of isoforms.
